# A Pipeline for Precise and Efficient Genome Editing by sgRNA-Cas9 RNPs in *Drosophila*

**DOI:** 10.1101/2020.05.07.080762

**Authors:** Kevin G. Nyberg, Joseph Q. Nguyen, Yong-Jae Kwon, Shelby Blythe, Greg J. Beitel, Richard W. Carthew

## Abstract

Genome editing via homology-directed repair (HDR) has made possible precise and deliberate modifications to gene sequences. CRISPR/Cas9-mediated HDR is the simplest means to carry this out. However, technical challenges remain to improve efficiency and broaden applicability to any genetic background of *Drosophila melanogaster* as well as to other *Drosophila* species. To address these issues, we developed a two-stage marker-assisted strategy in which embryos are injected with RNPs and pre-screened using T7EI. Using sgRNA in complex with recombinant Cas9 protein, we assayed each sgRNA for genome-cutting efficiency. We then conducted HDR using sgRNAs that efficiently cut target genes and the application of a transformation marker that generates RNAi against *eyes absent*. This allows for screening based on eye morphology rather than color. These new tools can be used to make a single change or a series of allelic substitutions in a region of interest, or to create additional genetic tools such as balancer chromosomes.

## INTRODUCTION

Genome editing by CRISPR/Cas9 has transformed research and development in the life and health sciences (1). Although use of CRISPR has expanded beyond the original genome editing capabilities (2–4), genome editing remains a very popular application. The Cas9 endonuclease associates with a single guide RNA (sgRNA), and the complex localizes to DNA sequences in the genome by following simple DNA:sgRNA base-pairing rules. The complex then introduces a double strand break (DSB) in the DNA, triggering repair of the broken ends. If there is available a separate DNA template that contains sequences homologous to the regions flanking the DSB, then homology-directed repair (HDR) can result in incorporation of the repair template into the genomic DNA. The repair template can be a homologous chromosome or an exogenous donor DNA molecule. Exogenous templates come in one of two forms: a single-stranded oligonucleotide or a double-stranded DNA plasmid. Plasmids can be much larger than oligonucleotides, allowing for modifications to be made at a greater distance from the DSB. In the absence of a repair template, non-homologous end joining (NHEJ) ligates the broken ends, resulting in stochastic insertions and deletions (indels) at the break site. Although NHEJ-mediated genome editing is useful for gene disruption, HDR affords precise and programmable alterations in genome sequence.

The adaptation of CRISPR/Cas9 editing to *Drosophila* research occurred shortly after its invention (5–7). The first-generation of methods used either expression plasmids or RNAs coding for Cas9 and sgRNA. Subsequently, homemade Cas9 protein complexed with tracrRNA and crRNA was shown to work in *Drosophila* (8). Additionally, a number of germline-specific Cas9 transgenic lines were generated (9–13), and these have been widely used by *Drosophila* researchers. Typically, a plasmid encoding a sgRNA driven by RNA polymerase III transcription is injected into *Cas9* transgenic embryos. This approach greatly increases the efficiency of germline editing events. It has been particularly beneficial for the development of CRISPR/Cas9-induced HDR in *Drosophila* (6, 10, 11, 14, 15).

CRISPR-induced HDR for genome editing is not straightforward because of two issues. First, computational design of a targeting sgRNA does not predict the efficiency of cleavage, which varies considerably at different target sites (16). This could be due to many reasons such as secondary structure within the sgRNA, stability of the sgRNA-DNA duplex, or accessibility of the target sequence within the context of chromatin. Therefore, editing of cultured cells often relies on multiple sgRNAs targeting one gene, as a way to “cover all bases” (16). This approach has also been developed for *Drosophila*, in which multiplexed sgRNAs are expressed from one vector (17). The second challenge for editing by HDR is that the individuals who have inherited the desired edits must be identified. This challenge exists because HDR resolution of DSBs is much rarer than NHEJ repair, and so the vast majority of individuals have not been edited. Furthermore, imprecise HDR can occur to create undesired edits. For *Drosophila*, molecular screening methods such as PCR and sequencing are time-consuming because G1 progeny of injected G0 flies must be individually assayed for precise HDR events.

As an alternative screening method, a visible transformation marker gene can be incorporated into the repair template plasmid (18). The marker gene is placed between the right and left homology arms used for template-driven repair. This affords rapid screening of G1 animals without their sacrifice. Use of such a selection scheme faces several challenges after transformants are identified. First, imprecise HDR events involving crossover repair at the site of a DSB are frequent occurrences (19). There, the repair template plasmid backbone is incorporated into the genome, and such events are scored positive using a transformation marker. To identify such events, HDR repair template plasmids can contain a *mini-white* gene in their backbones (https://flycrispr.org/; Addgene 80801), a modification inspired by recombinase-mediated cassette exchange (RMCE) vectors (20). The presence of *white+* in the plasmid backbone allows for a counter selection against the integration of the whole plasmid when incorporated into a *white* mutant background, ensuring that only the DNA between the homology arms integrates. However, the general utility of the *white* marker is limited by the necessity of using it in a *white^-^* genetic background and by the large size of its coding and control sequences. A second challenge is the presence of the marker gene at the site of editing. If the goal of editing is to determine the effects of precise base changes, then the marker gene must be removed prior to phenotypic analysis. Although ϕC31-mediated RMCE and FLP-FRT have been used to excise an HDR marker gene (15, 21), they leave scars in the form of ectopic sequences at the excision site. The PiggyBac transposase has been harnessed to excise a *3xP3-DsRed* marker gene from the edited site, and this approach has the benefit of leaving no sequence scar at the excision site (https://flycrispr.org/). An alternative scarless approach involves integration of the marker gene at the edited site, followed by a second round of HDR that replaces the marker gene with the desired edits (20, 22). Although reversion events are easily scored, the overall process requires two rounds of CRISPR/Cas9 injections and screening.

In summary, many developments have improved the scope and efficacy of genome editing in *Drosophila*. However, several impediments still remain until editing becomes as straightforward and efficient as more established genetic technologies in *Drosophila*. Here, we describe a series of modifications to the HDR-mediated editing procedure that overall enhance the success rate of achieving precise edits. Moreover, these enhancements can be adopted across a broad range of genetic backgrounds in *D. melanogaster* and even in other *Drosophila* species. We believe that this new procedure greatly expands the potential use for precise genome editing in *Drosophila*. The potential to modify one, two, or many more basepairs makes it a powerful tool for testing the phenotypic consequences of changes in genome sequence ranging from single base variants to more substantial differences.

## RESULTS

A survey of the existing technologies related to HDR-mediated editing by CRISPR identified several obstacles. These are listed in Figure 1. We systematically describe each obstacle and the method we developed to overcome it.

**Figure 1.**
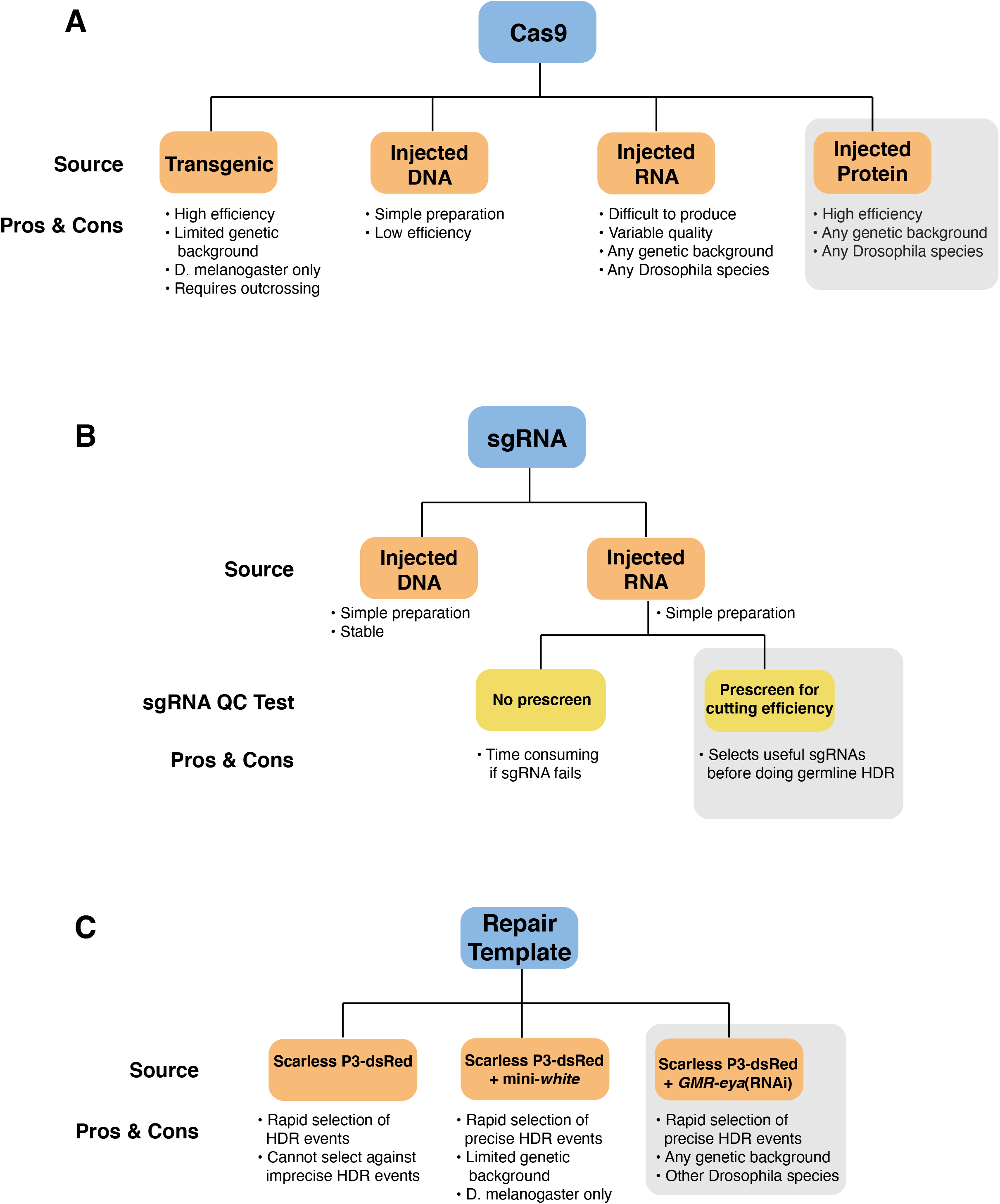
Sources of *Drosophila* CRISPR reagents and their individual pros and cons. (**A**) Sources of Cas9. Source shaded grey highlights the novel source used in this study. (**B**) Sources of sgRNA. Since some sgRNAs are inactive for inducing DSBs *in vivo*, a quality control (QC) test is preferable. Region shaded grey highlights the novel prescreening method to identify active sgRNAs. (**C**) Sources of donor plasmids to act as repair templates for HDR. This panel only shows plasmids related to the novel plasmid used in this study (shaded grey).

### Genome editing by RNP injection

The source of Cas9 used to induce DSBs varies, but the most commonly used source is a transgenic Cas9 specifically expressed in germ cells using regulatory sequences from the genes *vasa* or *nanos* (Fig. 1A). Although it usually induces DSBs with high efficiency, the reliance on a transgenic line limits the genetic background available for G0 founders and often complicates the background of G1 and subsequent generations. Cas9 plasmid and mRNA are less efficient and are more variable if injected. However, DSBs have been demonstrated to occur in cultured human cells using electroporation-mediated delivery of sgRNA and Cas9 in the form of *in vitro* assembled ribonucleoproteins (RNPs) (23, 24). This approach has also been adapted for generation of NHEJ-induced mutagenesis in *Drosophila* and *Caenorhabditis elegans* by microinjection (8, 25). However, the method for *Drosophila* relied upon homemade Cas9 protein combined with two RNAs rather than the standard sgRNA (8). Moreover, the efficiency of DSBs using RNPs was not directly compared to the use of *Cas9* transgenes. Finally, the utility of RNPs for HDR-mediated genome editing was not tested.

To determine the efficiency of RNPs for *Drosophila* editing, we assembled sgRNA-Cas9 RNPs by co-incubation of *in vitro*-synthesized sgRNA with commercially available recombinant Cas9 protein. The sgRNA was generated by T7 polymerase transcription of a synthetic DNA template as described (5). Preparation was as simple and reproducible as generating sgRNA-expression plasmids (Fig. 1B). Moreover, the commercial origin of the Cas9 protein makes it affordable and readily accessible to most *Drosophila* research groups.

An RNP solution designed to induce DSBs in the *forked* gene was injected into 328 *white* embryos using a previously validated sgRNA that targeted *forked* coding sequence (17). We then testcrossed the resulting 52 adults to *forked* mutants and scored G1 offspring for germline transmission of *forked* mutations induced by NHEJ. Of 39 crosses that produced G1 offspring, 23 (59.0%) resulted in one or more *forked* mutant offspring. We compared the efficiency of generating such mutants to the efficiency when the sgRNA alone was delivered into a transgenic *vasa-Cas9* line (BDSC #51324). Injection of 331 *vasa-Cas9* embryos resulted in 68 adults. When testcrossed to a *forked* mutant, 54 produced G1 offspring, and 12 (22.2%) of these crosses resulted in one or more *forked* mutant offspring. Thus, simple assembly and injection of RNPs was 2.5-fold more potent than using transgenic *vasa-Cas9* for inducing indel mutations in the *forked* gene (*p* = 0.0005, Fishers exact test). However, this *Cas9* transgene has slightly weaker efficiency than some other *Cas9* transgenes (11), and thus it will be interesting to see if RNPs using other sgRNAs also outperform this and other transgenes.

### sgRNA Screening

Clearly, use of RNPs expands the potential for inducing DSBs in any genetic background or even other *Drosophila* species. However, it has another important benefit. Given that sgRNAs with different target sites exhibit different endonuclease activities *in vivo*, prescreening the activity of various sgRNAs within the gene of interest would be advantageous before designing an appropriate repair template that fits with the selected sgRNA (Fig. 1B). At present, prescreening in *Drosophila* is not commonly performed because of the use of transgenic germline *Cas9* lines. Screening G1 or G2 animals constitutes a time-consuming and expensive process. However, injection of assembled RNPs induces DSBs not only in the germline but in the somatic cells of the animal as well. Therefore, we could screen for NHEJ-mediated mutations in G0 animals without having to generate HDR repair template plasmids.

The presence of indels induced by CRISPR can be detected by a T7 endonuclease I (T7EI) assay (Fig. 2A) (23). A prior study of RNP-mediated NHEJ mutagenesis in *Drosophila* found that the T7EI assay was able to detect indels in G0 and G1 adults (8). We reasoned that if such events could be detected in G0 embryos, then the time to analyze a sgRNA for cutting efficiency could be reduced from 12 - 25 days to 3 days. This has never been tested in G0 embryos. We injected a minimum of 8 embryos with RNPs, and purified genomic DNA from each individual after 24 hours. A small region surrounding the target site was amplified by PCR from individual DNA. If the RNPs had induced indels in a sizable number of an embryo’s cells, then the amplicons would be a composite of wildtype and mutant DNA duplexes. If the RNPs failed to induce many indels, then the amplicons would be primarily composed of wildtype strands. We then denatured the amplicons from each individual embryo source and hybridized the strands back together. This was followed by T7EI treatment. T7EI recognizes and cleaves mismatched heteroduplex DNA which arises from hybridization of wildtype and mutant DNA strands. If there were no mismatched heteroduplex DNAs, then the amplicons would remain at their original size, indicating that no indel mutations had been detectable in that individual embryo. However, cleavage of some amplicons by T7EI would indicate that a significant number of indel mutations had been induced in that individual embryo (Fig. 2A). We could then tally the fraction of injected embryos that scored positive in this T7EI assay for each tested sgRNA.

**Figure 2.**
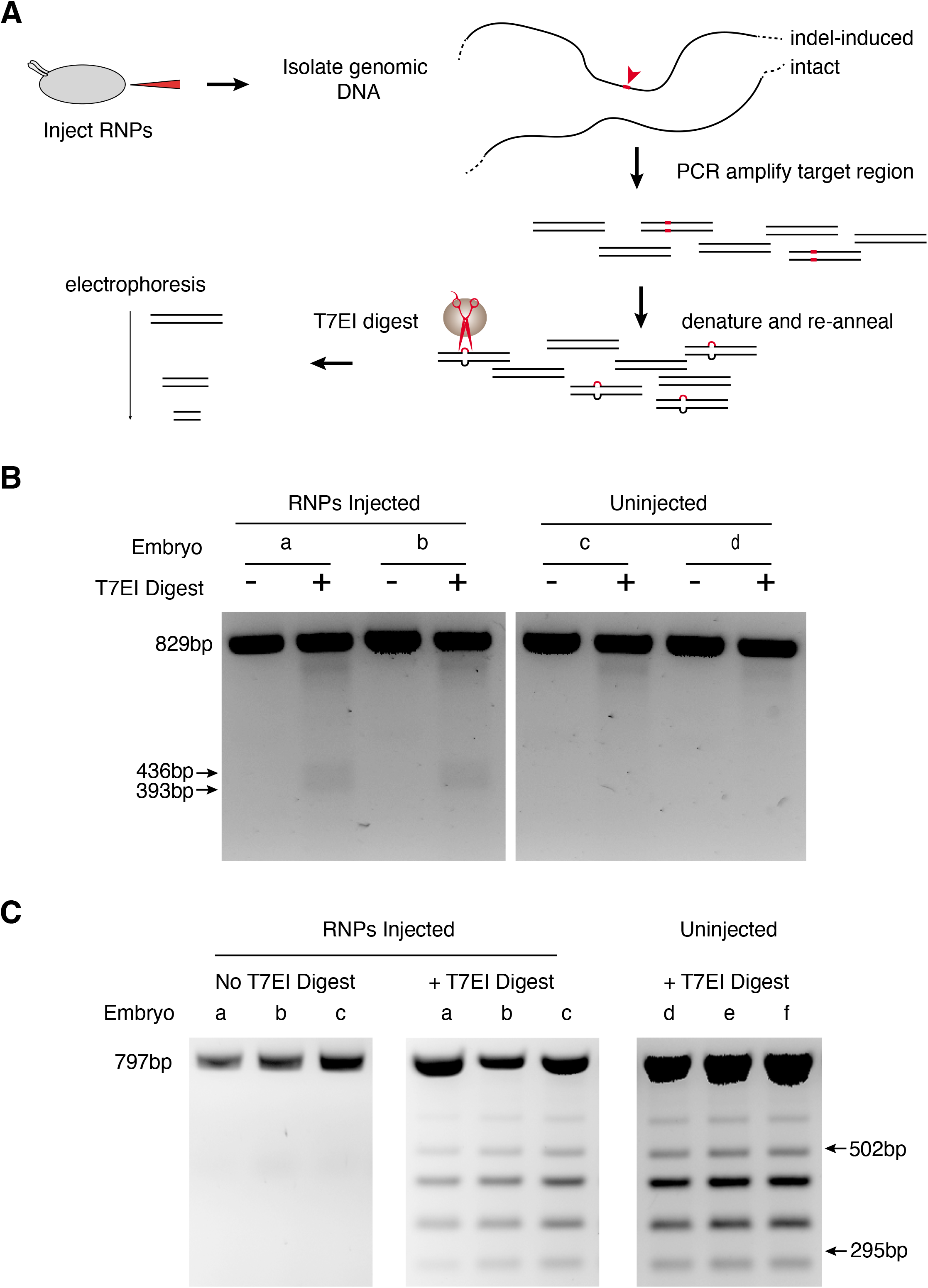
Screening sgRNAs for cleavage activity in vivo. (**A**) Schematic of the screening assay. Individual embryos are injected with RNPs composed of a particular sgRNA. Genomic DNA from each embryo is PCR-amplified, and amplicons are denatured and re-annealed. Heteroduplexes with mismatches due to indels in embryonic DNA are cleaved by T7EI enzyme. Gel electrophoresis identifies embryos with detectable cleavage events. (**B**) PCR products of a target site in the *forked* gene 892 bp in length were digested by T7EI as indicated. Shown are two representative embryos out of the nine assayed that were injected with *forked* RNPs. Also shown are two out of the six embryos that were uninjected. The predicted T7EI digest products are 393 and 436 bp (arrows). Although a minority of heteroduplexes derived from an embryo are T7EI-sensitive, they can be detected by this assay. A total of 6 out of 9 injected embryos showed evidence of T7EI sensitivity, whereas 0 out of 6 control embryos showed evidence. This difference is statistically signficant (*p*=0.0168, Fishers exact test). (**C**) A T7EI assay performed on a sgRNA that was inactive in vivo. The target region is located in non-coding DNA. Three of the 12 RNP-injected embryo samples are shown, and 3 of the 6 uninjected embryo samples are shown. Heteroduplexes from the uninjected samples show T7EI sensitivity that is likely due to inherent sequence heterogeneity in the strain at that particular locus. It is probably due to natural SNPs or Indels that are strongly heterozygous in the population that serves as the injected strain. The predicted T7EI digest products from NHEJ induced mismatches are 295 and 502 bp (arrows). Since the 502 bp band is obscured by a background band, the 295 bp band is diagnostic. Note that the 3 samples from RNP-injected embryos do not exhibit T7EI products of 295 bp size. Zero of 12 injected embryos showed evidence of having 295 bp products (*p* > 0.05, Fishers exact test).

We injected RNPs assembled from the *forked* sgRNA into syncytial embryos. T7EI reactions of amplicon heteroduplexes from 9 embryos were run on an agarose gel, revealing that 6 of the 9 samples produced cleavage products (Fig. 2B). T7EI treatment of PCR amplicons from uninjected embryos resulted in no cleavage products for the embryos tested. This difference in outcome depending on the sgRNA was statistically significant (*p*=0.0168, Fishers exact test). Thus, injection of RNPs into syncitial embryos is sufficient to induce NHEJ events that are frequent enough to be detected in late-stage embryos. We validated the method by testing 18 other sgRNAs targeting five different genes. For each sgRNA, we injected and analyzed a minimum of 8 single embryos and analyzed control uninjected embryos in parallel. The percentage of embryos with detectable cleavage events depended on the sgRNA, suggesting that sgRNA activity is quite variable. Of the total 19 sgRNAs tested, 5 sgRNAs (26%) failed to yield any embryos with detectable NHEJ events, and were statistically indistinguishable from control uninjected embryos (*p* > 0.05, Fishers exact test) (Fig. 2C). We confirmed that one of the failed sgRNAs was inactive for HDR-mediated editing by using it to induce DSBs in *Cas9* transgenic lines, accompanied by a repair template plasmid. Injection of ~900 G0 embryos failed to elicit a single HDR event. In contrast, five other sgRNAs that were positive in the T7EI assay and were then used for HDR-mediated genome editing, resulted in generating lines with the desired genome edits. From these experiments, we conclude that a significant fraction of sgRNAs are inactive in *Drosophila* despite being selected by computational prediction programs. These findings are consistent with studies in cell culture and mammals (16, 26).

### A broadly applicable repair template vector that incorporates a novel transformation marker

The injection of RNPs clearly provide benefits for rapid selection of active sgRNAs and for broader application to genetically diverse *Drosophila* backgrounds. However, the potentially broader applications are limited by the use of existing repair template vectors. The scarless 3xP3-DsRed donor vector should work across *Drosophila* and other insect genera owing to the fact that *3xP3-DsRed* can function in many insect species (27). However, the counter-selection marker *mini-white* gene only functions in *white* mutant genetic backgrounds. Thus, counter-selection for imprecise HDR events is not possible when using *mini-white* in other backgrounds or species with a wildtype *white* locus.

We have developed an alternative counter-selection marker to be used either independently or in conjunction with the scarless 3xP3-DsRed vector that will work in any genetic background and does not require the use of fluorescent microscopy to score. The counter-selection marker gene is composed of the GMR promoter driving a short hairpin RNA (shRNA) against the *eyes absent* (*eya*) gene, taken from the TRiP collection (HMS04515) (Fig. 3A). *Eya* is essential for proper compound eye development, and its loss results in small rough eyes (27). Since the GMR promoter is specifically active in compound eye cells, the *eya* gene should be knocked down by RNAi and generate small eyes. As proof of principle, we tested the effect of one copy of the *GMR-eya*(shRNA) marker on the adult eye phenotype and found that it produced a 100% penetrant small eyed phenotype. Notably, enough residual eye tissue is present in heterozygous flies to allow scoring of *white^+^* or 3XP3 fluorescent eye markers. We tested the general utility of this marker by constructing a new fourth chromosome balancer by insertion of *GMR-eya*(shRNA) into the *gat* gene on the fourth chromosome. This balancer, *GAT^eya^*, has a morphological phenotype more robust and easier to score than existing fourth chromosome balancers, and is also much healthier than the commonly used CiD balancer. Stocks of the new balancer have been deposited at the Bloomington Drosophila Stock Center (*GAT^eya^/CiD*, BDSC #90852; *GAT^eya^/Crk^dsRed^*, BDSC #90851).

**Figure 3.**
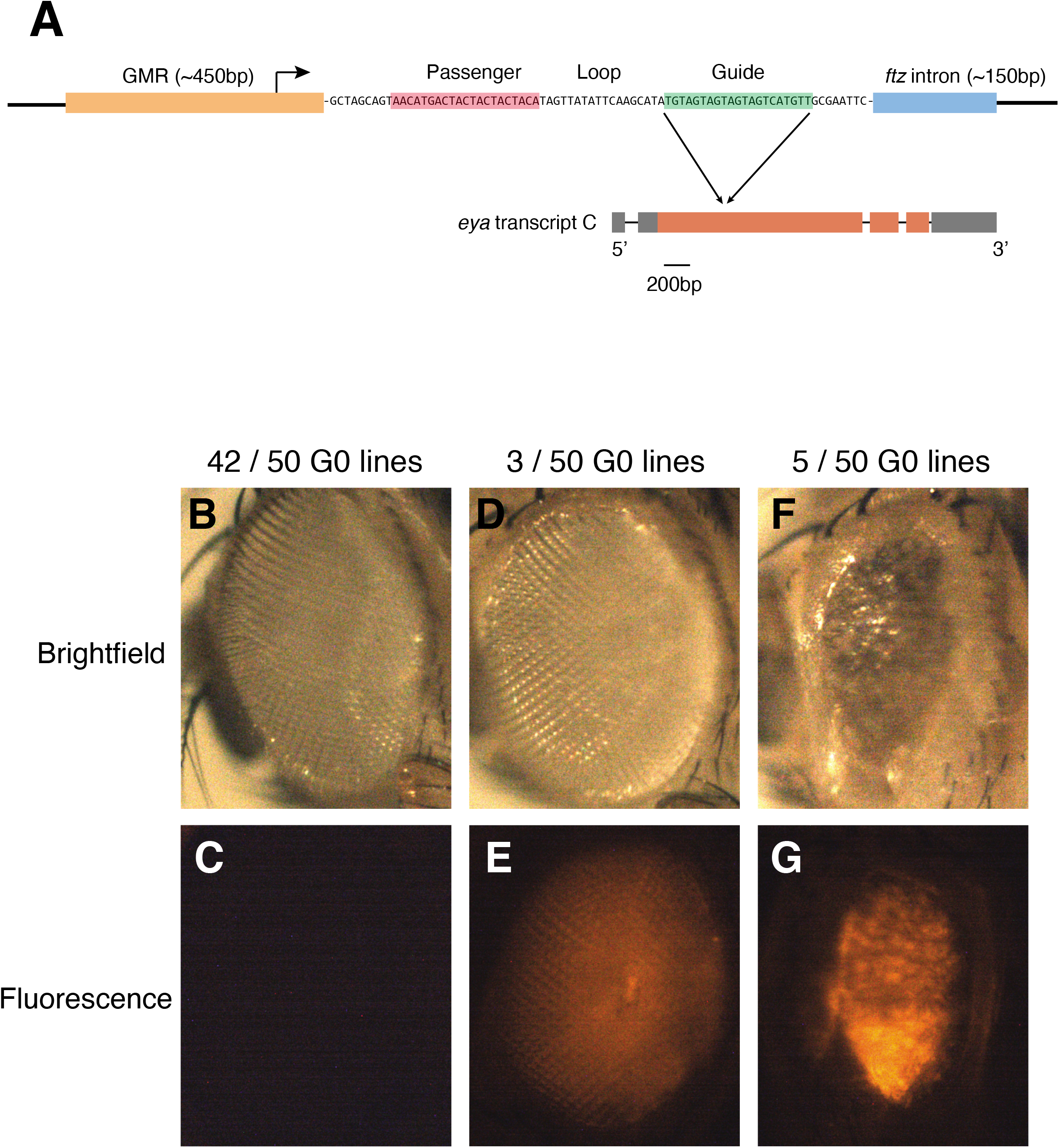
The modified plasmid backbone for HDR editing. (**A**) Shown is the transgenic marker for counterselection of imprecise HDR events. The GMR element contains 5 tandem binding sites for the transcription factor Glass fused to the Hsp70 minimal promoter. The transcript contains a shRNA stem-loop followed by an intron from the *ftz* gene to facilitate transcript stability. After the shRNA is processed by Drosha and Dicer, the guide RNA strand is loaded into RISC. The guide RNA is perfectly complementary to all mRNA isoforms of *eya*. Shown only is isoform C, and the location of the RNAi target is indicated. (**B-F**) Eye phenotypes of adults that had been injected with RNPs and the *forked* HDR donor plasmid. **(B,D,F)** Eyes visualized with brightfield illumination. (**C,E,G**) Same eyes visualized for DsRed fluorescence. All eyes are oriented anterior left and dorsal top. Listed above panels are the number of G0 lines that produced G1s with the same phenotypes as the ones shown. (**B,C**) Adult without apparent HDR event. (**D,E**) Adult with DsRed expression and no *eya* RNAi phenotype. (**F,G**) Adult with DsRed expression and an *eya* RNAi phenotype.

To construct a counter selectable repair template backbone, the *GMR-eya*(shRNA) marker was inserted into pBlueScript to create the pBS-*GMR-eya*(shRNA) plasmid (Addgene #157991; Drosophila Genomic Resource Center #**). Upon linearization at its multi-cloning site, typically with *EcoRV*, the pBS-*GMR-eya*(shRNA) plasmid serves as the backbone for construction of a donor plasmid. Using Gibson assembly, the scarless *3xP3-DsRed* transformation marker flanked by 1 kb homology arms and desired genomic sequence modification can be assembled with the *pBS-GMR-eya*(shRNA) backbone. Thus, precise HDR events can be discriminated from integration of the entire plasmid into the genome by the presence of DsRed fluorescence and the absence of a small eye phenotype. An additional benefit to replacing the mini-*white* gene with *GMR-eya*(shRNA) is that *mini-white*, whose size belies its name, is ~2,800 bp while *GMR-eya*(shRNA) is only 820 bp. This reduces the donor plasmid size, making it easier to construct repair templates.

We tested the efficacy of the pBS-*GMR-eya*(shRNA) vector by placing left and right homology arms targeting the *forked* gene on either side of 3xP3-DsRed. The HDR donor plasmid was designed to induce insertion of 3xP3-DsRed into the coding region of *forked*, thus disrupting gene function. We injected 461 *white* embryos with the HDR donor plasmid and Cas9 protein complexed with the *forked* sgRNA used previously. Of the 50 fertile G0 adults that emerged after injection, three produced G1 offspring that expressed DsRed in eyes that were normal sized (Fig. 3B-E), while five produced G1 offspring that expressed DsRed in small *eya*-like eyes (Fig. 3F,G). Normal sized eyes with DsRed fluorescence in the G1 offspring indicate precise HDR of the target gene. Nine G1 adults met these criteria. Another nine G1 adults displayed small eyes that expressed DsRed, presumably the result of imprecise HDR events. G1 adults with normal-sized DsRed-expressing eyes were testcrossed to a *forked* stock, and six crosses generated 100% penetrant *forked* mutant phenotypes. Sequence analysis showed repair with both homology regions matching the donor plasmid sequence, confirming that we had precisely introduced the DsRed marker in the six lines.

As another test, we targeted the essential gene *crk* on the fourth chromosome. We established a DsRed positive, non-small eyed line in which the *crk* coding sequence was replaced by the *3xP3-DsRed* marker, thereby creating the first fluorescently-marked fourth chromosome balancer that we are aware of (*Crk^dsRed^*, BDSC#90850 and #90851).

Other loci were independently targeted with donor plasmids and sgRNAs, and in four out of the five targeted, some percentage of DsRed-positive G1 adults had small *eya*-like eyes (Table 1). The percent ranged from 9 to 49% depending on the target. Overall, including results from the *forked* editing, an average of 29% of edited germline events were imprecise. Thus, pBS-*GMR-eya*(shRNA) is an effective and useful counter-selection marker for HDR-mediated editing.

**Table 1.** Scoring for precise HDR events using *eya*(shRNA) counterselection.

| HDR Construct | Number injected | Number fertile G0 adults | Number G1 DsRed+ *eya*+ | Number G1 DsRed+ *eya*(shRNA) |
| --- | --- | --- | --- | --- |
| 1 | 342 | 84 | 52 | 5 |
| 2 | 340 | 70 | 27 | 26 |
| 3 | 344 | 95 | 12 | 4 |
| 4 | 295 | 52 | 18 | 0 |
| 5 | 310 | 64 | 14 | 10 |

## DISCUSSION

We have developed tools and protocols to implement a two-step genome editing workflow suitable for making precise changes at targeted sites in *Drosophila* (Fig. 4). Detailed protocols for each step in the workflow are available in the Supplementary Information, and plasmid reagents have been deposited in Addgene and the Drosophila Genome Resource Center (DGRC). Our workflow adds to the available methods that do not leave scars in the genome, such as those that occur when ablating PAM sites or using integrase-mediated excision to remove selectable markers. Injection of Cas9-sgRNA molecular complexes is several fold more efficient at generating DSBs than *vasa-Cas9*. Injection of RNPs is also well-suited for making changes in other *Drosophila* species. This feature realizes the potential for genome editing to make changes at a gene’s native locus and in its native genome. For example, the function of sequences that have diverged between two *Drosophila* species can now be tested in their native context rather than in *D. melanogaster*, as had been previously done.

**Figure 4.**
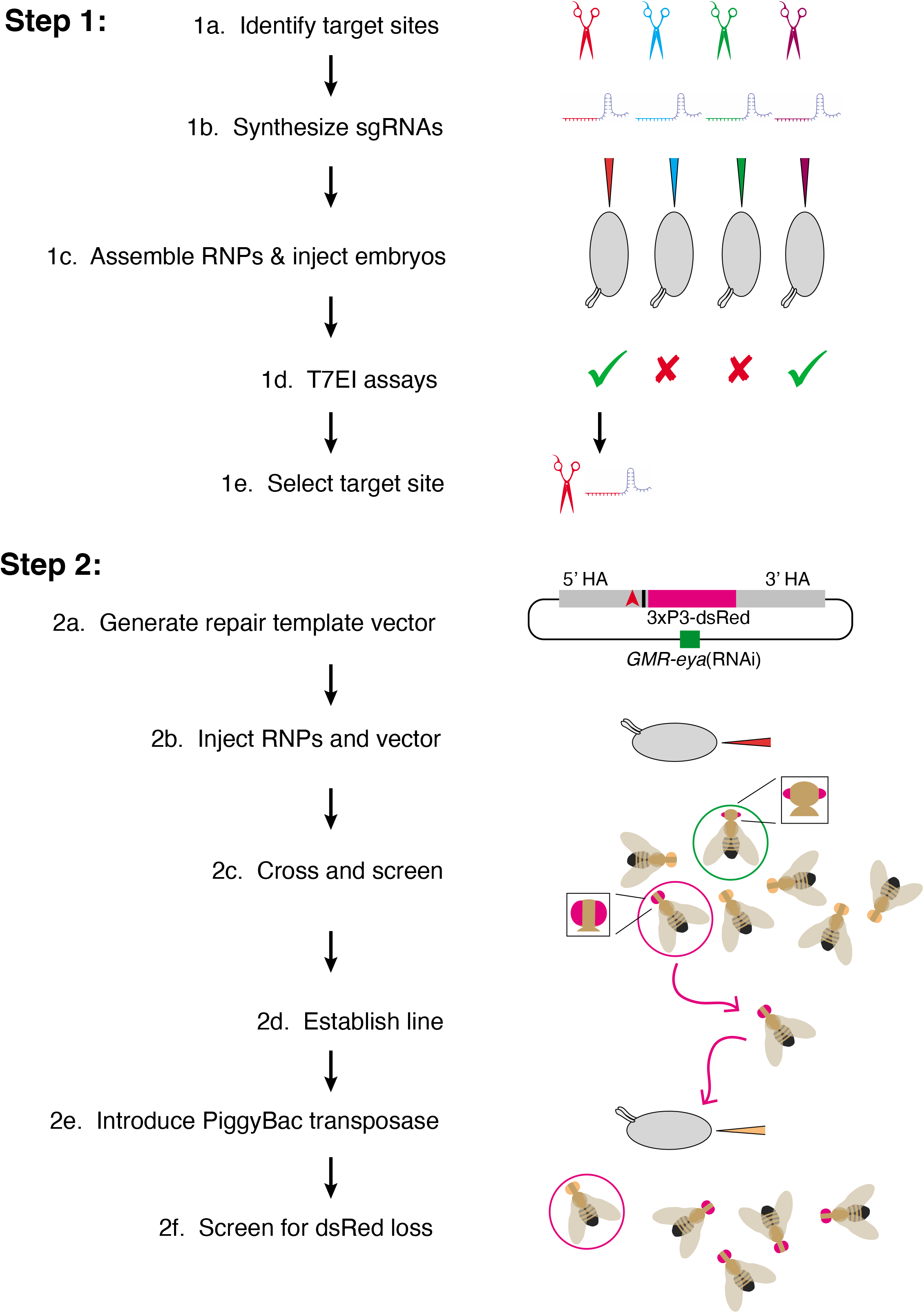
Workflow for two-step genome editing. (**1a**) Target sites flanking the area to be edited are identified (red, blue, geen, purple) using online tools searching for optimal targets and with minimal off-target cleavage. (**1b**) Sequences from the selected target sites are transcribed *in vitro* to generate sgRNAs. (**1c**) Cas9 protein is incubated with sgRNAs before injection into embryos. (**1d**) Active sgRNAs that cleave embryo DNA are identified by T7 endonuclease I reactions. (**1e**) One of the active sgRNAs is chosen for genome editing in Step 2. (**2a**) Homology arms flanking the region of interest are cloned into the pBS-*GMR-eya*(shRNA) donor plasmid. In this example, the CRISPR target site (red triangle) is 5’ to the bases to be edited (black bar). (**2b**) Embryos are injected with the repair template plasmid and RNPs composed of sgRNA and Cas9 protein. (**2c**) Adult flies that develop from injected embryos are crossed back to the parental line. G1 progeny are screened for the DsRed marker. Positive G1 animals may have small eyes due to *eya*(shRNA) but these are not selected (green circle). Only positive G1 animals with normal eyes are selected (red circle). (**2d**) These are crossed to make purebred lines and molecularly analyzed to determine if they contain the desired editing events. (**2e**) PiggyBac transposase is expressed in the germline, either by a single cross to a transgenic line, or in this example, by embryo injection of a plasmid expressing the transposase. (**2f**) Since the DsRed marker is dominant, adult flies developing from injected embryos that do not have red fluorescent eyes are then crossed and analyzed with molecular tests to determine whether they have precisely excised the marker gene. Only the intended genomic edit remains.

The first step in the workflow is the design and testing of candidate sgRNAs. Since sgRNAs vary in their ability to induce DSBs in genomes, it is worthwhile to test a number of sgRNAs for making the desired edit. We found that 26% of tested sgRNAs are inactive in *Drosophila* embryos. Therefore the first step in the workflow is to rapidly synthesize a few sgRNAs by *in vitro* transcription, followed by assembly of RNPs with Cas9 protein. A simple PCR-based endonuclease assay then measures the ability of each sgRNA to induce NHEJ indel mutations in injected embryos. This procedure takes at most 3 days and will work in other species besides *D. melanogaster*. The 3 days effort potentially saves 2 to 3 months of wasted effort to perform HDR using a sgRNA that does not work *in vivo*. Once a sgRNA is selected, the second step in the workflow is taken. Homology arms, desired genome alterations, and the scarless 3xP3-DsRed marker are Gibson assembled with pBS-*GMR-eya*(shRNA) such that the arms flank the scarless DsRed marker. This construct is co-injected with matching RNPs into the strain or species of choice, and G1 individuals with red fluorescent eyes are selected. Although it is easier to screen such individuals in a *white* mutant background, it is not necessary. Red fluorescence can be detected in the adult eyes of *white^+^ D. melanogaster* expressing *3xP3-DsRed*, although fluorescence is only visible in a small spot of 10 ommatidia (data not shown and (27)). This property of *3xP3-DsRed* fluorescence is also observed in other *Drosophila* species (28). If compound eye fluorescence is too weak, the adult ocelli or larval Bolwigs organs can also be examined (27, 28).

Our method uses a novel marker placed in a pBluescript backbone to select against editing events in which the entire vector has integrated into the genome. The marker uses RNAi to knock down expression of the endogenous *eya* gene, resulting in a small eye. The *eya* shRNA is designed to work in any *D. melanogaster* strain and will work in the closely related *D. erecta* species. *D. simulans, yakuba*, and *ananassae* have only single base variant in the sequence targeted by RNAi, so the backbone can easily be modified via site-directed mutagenesis for these species. Since we have found that 29% of DsRed transformants also have *eya*(RNAi) phenotypes, our method reduces the laborious molecular characterization of false-positives.

There are added potential functions for the *GMR-eya*(shRNA) gene in that its insertion into any chromosome renders a simple dominant phenotype that is highly penetrant. Thus, it can be used to mark any balancer chromosome. We have done so for the fourth chromosome balancer, and this new balancer (*GAT^eya^*) is deposited at the Bloomington Drosophila Stock Center. It should be possible to use CRISPR/HDR to insert *GMR-eya*(shRNA) in other balancers with subtle dominant markers, such as *TM6,Ubx*.

The final step in the workflow is the scarless excision of *3xP3-DsRed* from the genome by PiggyBac-mediated transposition. Scarless editing requires the presence of a TTAA motif at the target locus, which adds some restriction to target selection. However, it circumvents the need for a second round of injections because *D. melanogaster* carrying the marker can be crossed to existing transgenic lines that express the PiggyBac transposase (29). To apply *3xP3-DsRed* excision to species other than *D. melanogaster*, a subsequent injection step is needed to introduce the PiggyBac transposase. There are appropriate expression plasmids that are freely available (30).

In conclusion, the applicability of our method to many types of experiments in a wide variety of genetic backgrounds makes it a valuable addition to the existing methods and tools for scarless genome editing available to the *Drosophila* research community.

## MATERIALS AND METHODS

A detailed protocol and description of reagents is provided in Supplementary Information. Statistics were performed using Graphpad software.

### *Drosophila* Strains

*Drosophila* were raised on standard cornmeal-molasses food at room temperature. The *vasa-Cas9* strain (BDSC#51324) used for some experiments is from (10). This transgene is located on the third chromosome and has been shown to have slightly weaker efficiency than some other Cas9 transgenes (11). Unless otherwise stated, all injections of CRISPR RNPs, sgRNAs, and donor plasmids were into a *w^1118^* strain.

### Plasmids

pBS-GMR-*eya*(shRNA) is derived from pBSII-KS(-), and contains a marker gene GMR-*eya*(shRNA), which is comprised of the <u>G</u>lass <u>M</u>ultimer <u>R</u>eporter (GMR) enhancer driving an *eya*(shRNA) from the Transgenic RNAi Project (TRiP) (31). GMR is a synthetic eye-specific enhancer composed of five tandem repeats of a 29 bp element from the *Rh1* gene (32). Tests using *GMR-Gal4* to drive UAS-*eya*(shRNA) from the TRiP lines HMS04515 and HM05716 revealed that HMS04515 produced a stronger eye phenotype than HM05716. Therefore, the *eya*(shRNA) sequence from HMS04515 was used to construct the GMR-*eya*(shRNA) marker gene. This shRNA targets all three transcripts of the *eya* gene. A PCR fragment containing the GMR enhancer and the *hsp70* minimal promoter was amplified from pGMR DNA (33) with *XhoI* and *XbaI* ends. This fragment was inserted into a version of pKanC5 (34) in which a linker containing *XhoI* and *AvrII* sites had been inserted between the *BbvI* and *AscI* sites. The GMR cassette was then shuttled into the HMS04515 Valium20 *eya*(shRNA) plasmid, inserted between the *StuI* and *XbaI* sites using Gibson assembly (35). This replaced the UAS enhancer and upstream Gypsy insulator with the GMR cassette. The ~440 bp Gypsy insulator was eliminated to minimize the final size of the gene. The efficacy of the resulting GMR-*eya*(shRNA) Valium construct was tested by integrating it into the genome by ϕC31-mediated recombination (36). Once validated, the GMR-*eya*(shRNA) cassette was amplified by PCR from the Valium plasmid, and inserted into the *KpnI* site of pBSII-KS(-) using Gibson assembly (34). The resulting pBS-GMR-*eya*(shRNA) plasmid has been deposited with AddGene (#157991) and the Drosophila Genomics Resource Center (#****). Its annotated sequence is found in Supplementary File 1.

To construct a HDR donor plasmid, 3xP3-DsRed and homology arms flanking the targeted region of interest were PCR amplified to generate overlapping regions of homology for cloning into pBS-GMR-*eya*(shRNA) via Gibson Assembly (35). The 3xP3-DsRed marker gene from the pScarlessHD-DsRed plasmid (Addgene #64703) was PCR amplified. It is flanked by piggyBac TTAA transposition sites that can be used to cleanly remove the entire marker gene after successful integration of the modification via HDR. The ~1 kb 5’ homology arm was PCR amplified from *D. melanogaster* with a region homologous to the pBS-GMR-*eya*(shRNA) vector on the 5’ end. The ~1 kb 3’ homology arm was amplified from *D. melanogaster* with a region homologous to the pBS-GMR-*eya*(shRNA) vector on the 3’ end. The PCR amplicons were appended with sequence corresponding to sequence at a native TTAA site in the genome near the sgRNA site (ideally less than 30 bp) or within a TTAA site in the intended modification. A DNA fragment encompassing the sequence modification of interest was synthesized using gBlocks (IDT) with regions appended on both ends to facilitate Gibson assembly. These DNA fragments, the amplicons, and the pBS-GMR-*eya*(shRNA) vector linearlzed with *EcoRV*, were altogether assembled using New England Biolabs (NEB) NEBuilder HiFi DNA Master Mix. This generated the HDR donor plasmid. Inserts into the *EcoRV* site of pBS-GMR-*eya*(shRNA) can be sequenced using the custom primer 5’-actgggctcgaggcgatc-3’ on one side of the *EcoRV* site and custom primer 5’-ggcggccgctctagaactag-3’ or the standard M13-forward and T7 primers on the other side.

The *forked* HDR donor plasmid was composed of the following elements:

1. 5’ homology arm: X:17,269,009 to X:17,270,001 (Flybase release 6)
2. 3xP3-DsRed cassette: 5’-TTAA-(3xP3-DsRed)-TTAA-3’
3. *forked* mutagenesis fragment: This fragment of *forked* (X:17,270,002 to X:17,270,050) spanned from a TTAA site where 3xP3-DsRed was ligated to the sgRNA cut site. We had to introduce several synonymous mutations in the targeting fragment. There were no nearby TTAA sites, and so we created one with a synonymous mutation that changed a TCAA site (X:17,270,002 to X:17,270,005) in the genome to TTAA. We inactivated the sgRNA cut site in the fragment (X:17,270,034 to 17,270,050) with 2 synonymous mutations, as is common practice. The novel TTAA site is 45 bp from the sgRNA cut site. The fragment sequence with the synonymous mutations underlined is:

5’-T<u>T</u>AAGTTTCTGGTGCTCGAGGCCGGCGGCTCTTTGTACGTCCGTGC<u>T</u>CG<u>T</u>-3’
4. 3 ‘homology arm: X:17,270,051 to X:17,271,035 (FlyBase release 6)

### sgRNA and RNP Preparations

Candidate sgRNAs were identified using flyCRISPR Optimal Target Finder (http://targetfinder.flycrispr.neuro.brown.edu). High stringency filtering was used, and only NGG PAM sites were considered. Potential off-target sites were minimized to 0 predicted off-target sites. The sgRNA target site sequence was validated in the particular *Drosophila* strain being injected. This was done by Sanger sequencing the site from the strain’s genomic DNA.

The sgRNAs were synthesized by i*n vitro* transcription using T7 RNA polymerase. Transcription templates were created by PCR using a high fidelity polymerase. PCR amplification used the plasmid pU6-BbsI-chiRNA as template (6). pU6-BbsI-chiRNA was a gift from M. Harrison, K. O’Connor-Giles and J. Wildonger (RRID:Addgene_45946). The forward primer contained the T7 promoter sequence at the 5’ end followed by the sgRNA sequence (without the PAM) and then 19 bases complementary to the plasmid at the 3’ end. The complementary site on the plasmid corresponds to the 5’ end of the sgRNA scaffold at position 590.

5’-TTAATACGACTCACTATAGG[sgRNA_sequence]GTTTTAGAGCTAGAAATAG-3’

The reverse primer was a 20 base oligonucleotide complementary to the 3’ end of the sgRNA scaffold ending at position 669.

5’-AAAAGCACCGACTCGGTGCC-3’

The PCR amplicon was used in a MEGAscript in vitro transcription reaction (ThermoFisher #AM1333) supplemented with 0.5 μL Ribolock RNase inhibitor (ThermoFisher #EO0381) at 37°C overnight. The resulting sgRNA was purified with a Monarch RNA Purification Kit (NEB).

sgRNA-Cas9 RNPs were prepared fresh immediately before injecting embryos. 2.36 μg sgRNA was incubated with 11.9 μg Cas9-NLS recombinant protein (IDT #1081058) in 5 μL of 150 mM KCl. They were incubated at room temperature for 10 min followed by centrifugation at maximum speed for 10 min at room temperature. Supernatant was then loaded into injection needles.

The sequence of the sgRNA synthesized to target the *forked* gene was: 5’-UUGUACGUCCGUGCACGCGA-3’. It corresponds to X:17,270,034 to 17,270,056 (includes the PAM site)

### *Drosophila* Embryo Injections

Injections were performed in pre-cellularized embryos without dechorionation using Gompel and Schröder’s method (http://gompel.org/wp-content/uploads/2015/12/Drosophila-transformation-with-chorion.pdf). After injection, any embryos that were skipped during injection due to age or other defects were ruptured with a needle. Then, as much halocarbon oil was removed from the coverslip holding the embryos as possible. For T7EI assays, the coverslip with embryos was placed on an egg-laying plate, and the plate was incubated in a humid chamber at 25°C for 24 hours. For HDR editing, the coverslip with embryos was placed in a standard fly vial.

For injections to induce NHEJ, only freshly-prepared sgRNA-Cas9 RNPs were injected. To induce HDR edits, RNPs were assembled as described above except donor plasmid DNA was also added to the reaction. 2.36 μg sgRNA was incubated with 11.9 μg Cas9-NLS recombinant protein (IDT #1081058) plus 0.6 pmoles plasmid DNA in 5 μL of 150 mM KCl. They were incubated at room temperature for 10 min followed by centrifugation at maximum speed for 10 min at room temperature. Supernatant was then loaded into injection needles.

### Assay for sgRNA-Cas9 Mediated DNA Cleavage

Injected embryos were individually extracted using a single-fly genomic prep (https://kumarlab.bio.indiana.edu/7_labresources/protocols/016%20Single%20Fly%20Genomic%20DNA%20Extraction.pdf). L1 larvae or late-stage embryos only were chosen, indicating survival of the injection process. 3-5 μL genomic DNA was used as a template for a 50 μL PCR reaction in which primers were used that bounded the CRISPR target site being assayed. The amplicon was designed to be ~700 - 1200 bp in length with the target site located close to the center of the amplicon. The PCR product from one embryo was heat-denatured and slowly allowed to reanneal. 10 μL re-annealed PCR product was digested with 2 units T7 Endonuclease I (NEB #M0302L) in NEBuffer 2 for 60 min at 37°C. Reaction products were run on a 2% (w/v) agarose gel with ethidium bromide and 0.5X TBE. Samples with cleavage products at expected sizes often showed cleavage bands that were very faint.

### Genetic Screening

To screen for the presence of the GMR-*eya*(shRNA) marker, we examined G1 adults under standard dissecting microscopes for presence of small rough eyes. Such animals were annotated and then discarded. To screen G1 adults for the DsRed marker, we used a Nikon SMZ 1500 stereoscope equipped with a Fluorescence Illuminator. DsRed expression from the reporter gene used in this study becomes easily detectable in white eyes of adults. If the eyes have normal pigmentation, then the DsRed fluorescence is difficult to visualize except for a spot of ~10 ommatidia within the overall eye. Positive G1 adults were crossed to an appropriate balancer line. From these balancer crosses, siblings with both the DsRed phenotype and the balancer phenotype were crossed to form homozygous or balanced lines.

To precisely excise the DsRed marker, lines were crossed to a stock expressing the piggyBac transposase. The piggyBac transposase transgene is under control of the *α-tubulin* promoter and is tightly linked to a 3XP3-CFP transgenic marker (37). This is located on chromosome 2 (Bloomington Stock Center #32070). The stock also contains 3rd chromosome balancers (MKRS/TM6B,Tb), facilitating tracking of the 3rd chromosomes independent of the piggyBac transposase. Heterozygous offspring were crossed to appropriate balancer strains, and their offspring were screened for absence of DsRed and CFP fluorescence in adult eyes. Excision of DsRed occurs about 10% of the time. Positive animals were again crossed to a balancer strain to establish balanced stocks.

### Molecular screening

To confirm integration of the Ds-Red marker gene into the correct genomic location, we used a PCR reaction that amplifies DNA sequence from within the reporter gene sequence to outside the homology region on both the 5’ and 3’ sides of the reporter gene. The amplicons from these PCR reactions were Sanger sequenced to confirm scarless repair at both the target sites and throughout both homology regions. To screen for correct HDR after the DsRed excision, the entire edited locus was amplified via PCR using primers outside the homology regions. The amplicon was Sanger sequenced to confirm the presence of expected sequence edits. All diagnostic PCRs were performed using genomic DNA extracted from single flies following the squish prep protocol (https://kumarlab.bio.indiana.edu/7_labresources/protocols/016%20Single%20Fly%20Genomic%20DNA%20Extraction.pdf).

### *Gat^eya^* and *Crk^dsRed^* Fourth Chromosome Balancers

The *Gat^eya^* balancer fourth chromosome was created by targeting the *Gat* locus using the two sgRNAs detailed below and an HDR template constructed by Gibson assembly of 3xP3-DsRed flanked by *Gat* 5’ and 3 ‘homology arms in the pBS-*GMR-eya*(shRNA) backbone. Homologous repair from this template should result in deletion of all but the first 10 amino acids of the *Gat* open reading frame. Of three DsRed positive lines established from *vasa-Cas9* G0 animals (BDSC stock #55821), two were precise repair events (*Eya+* eyes) and one was an integration event (*eya* eyes). An *eya* line was established from the G0 with the integration event. This line was then crossed to the Piggybac transposase line to excise the DsRed transformation marker. A single F1 male lacking 3xP3-DsRed was used to establish the *Gat^eya^* chromosome. Neither the nature of the original insertion event nor the molecular nature of the *Gat^eya^* mutation produced by the dsRed excision has been investigated. The *vasa-Cas9*, piggyBac transposase, other markers and potential off-target mutations were removed from the background by backcrossing the *Gat^eya^* chromosome against *w^1118^* for five generations. The *Gat^eya^* chromosome serves as a fourth chromosome balancer because there is essentially no recombination on the fourth chromosome, the *Gat^eya^* mutation causes a dominant small-eye phenotype, and it is recessive lethal (1,435 out of 1,435 adults in a stock of *Gat^eya^/Crk^dsRed^* were DsRed+, *eya* RNAi).

*The a*ssembled order of the circular *Gat*/Scarless dsRed HDR template construct was: *pBS eya*(*shRNA*) backbone…*Gat* 5 ‘homology arm…3xP3-DsRed…*Gat* 3 ‘homology arm…*pBS eya*(*shRNA*) backbone. The sequences of the primers used to create that *Gat* targeting construct were (*Gat* sequences underlined):

Gat 5 ‘homology arm forward: 5 – ‘CCGGGCTGCAGGAATTCGAT<u>CAGGATCAATAGCCAAGTCGATCT</u> – 3’ Gat 5 ‘homology arm reverse: 5 – ‘CTTTAACGTACGTCACAATATGATTATCTTTCTAGGG<u>TTAAGTCACCATCGCTTGCGGA</u> – 3’
Gat 3 ‘homology arm forward: 5 – ‘GAGCAATATTTCAAGAATGCATGCGTCAATTTTACGCAGACTATCTTTCTAGGG<u>TTAAAGTGGTATGCCAGAAATATCTAG</u> – 3’
Gat 3 ‘homology arm reverse: 5 – ‘CGACGGTATCGATAAGCTTGAT<u>CATATTCACTCTTGTGAATAGACAC</u> – 3’

To construct *Gat* sgRNA plasmids, the following oligonucleotides were annealed and ligated into pU6-BbsI-chiRNA (RRID:Addgene_45946) to create two plasmids that produce sgRNAs targeting the 5 ‘and 3 ‘regions of the *Gat* gene:

*Gat* sgRNA1 forward: 5’-CTTCGCCGCAAGCGATGGTGACGG-3’
*Gat* sgRNA1 reverse: 5’-AAACCCGTCACCATCGCTTGCGGC-3’

*Gat* sgRNA2 forward 5’-CTTCGTTGTCGTACTTACTTAAAG-3’
*Gat* sgRNA2 Reverse 5’-AAACCTTTAAGTAAGTACGACAAC-3’

The *Crk^dsRed^* fourth chromosome balancer was created by targeting the *Crk* locus with an HDR template composed of the 3xP3-DsRed marker flanked by *Crk* 5 ‘and 3 ‘homology arms (see below) in the pBS-*GMR-eya*(shRNA) backbone. Homologous repair from the HDR template should result in the replacement of almost the entire open reading frame region of the *Crk* gene with the DsRed gene, from 48 bp 5 ‘of the start codon to 121 bp 5 ‘of the TAA stop codon. Two DsRed *Eya+* G1’s were identified from *vasa-Cas9* G0 animals. One line was established from a single G1 male to found the *Crk^dsRed^* chromosome. The precise nature of the repair event that created the *Crk^dsRed^* mutation has not been investigated. The *vasa-Cas9* and potential off-target mutations were removed from the background by backcrossing the *Crk^dsRed^* chromosome against *w^1118^* for five generations. The *Crk^dsRed^* chromosome serves as a fourth chromosome balancer because there is essentially no recombination on the fourth chromosome, the *Crk^dsRed^* mutation causes a dominant eye phenotype, and it is recessive lethal.

The assembled order of the HDR template construct is: *pBS eya*(*shRNA*) backbone…*Crk* 5’ homology arm…3xP3-DsRed…*Crk* 3 ‘homology arm…*pBS eya*(*shRNA*) backbone. The sequences of the primers used to create the *Crk* targeting construct were (*Crk* sequences underlined):

Crk 5 ‘homology arm forward: 5 – ‘CCGGGCTGCAGGAATTCGAT<u>ATTTTTGATCCTAGCTTCAAAATCT</u> – 3 ’
Crk 5‘ homology arm reverse: 5 –‘ CTTTAACGTACGTCACAATATGATTATCTTTCTAGGG<u>ATAAATAGAAATTATGTGATATAATGCAAATATA</u> – 3’
Crk 3‘ homology arm forward: 5 –‘ GAGCAATATTTCAAGAATGCATGCGTCAATTTTACGCAGACTATCTTTCTAGGG<u>AATTGGAAATAGGTGACATTATTAAAGTCA</u> – 3’
Crk 3‘ homology arm reverse: 5 –‘ CGACGGTATCGATAAGCTTGAT<u>AGAAGCACTAACTAACTATTGATCTAAAGAT</u> – 3’

To construct *Crk* sgRNA plasmids, the following oligonucleotides were annealed and ligated into pU6-BbsI-chiRNA targeting the 5‘and 3‘ regions of the *Crk* gene:

*Crk* sgRNA1 forward: 5’-CTTCGAATTTCTATTTATTTAATC-3’
*Crk* sgRNA1 reverse: 5’-AAACGATTAAATAAATAGAAATTC-3’

*Crk* sgRNA2 forward 5’-CTTCGGATAAGACTGCATTAAAAT-3’
*Crk* sgRNA2 Reverse 5’-AAACATTTTAATGCAGTCTTATCC-3’

## ACKNOWLEDGEMENTS

Fly stocks from the Bloomington Drosophila Stock Center are gratefully appreciated. Plasmids were from the Drosophila Genomics Resource Center. Financial support was provided from the NIH (F32GM122349, K.G.N.; R01GM108964, G.J.B.; R35GM118144, R.W.C.).

---

**Supplementary File 1.**
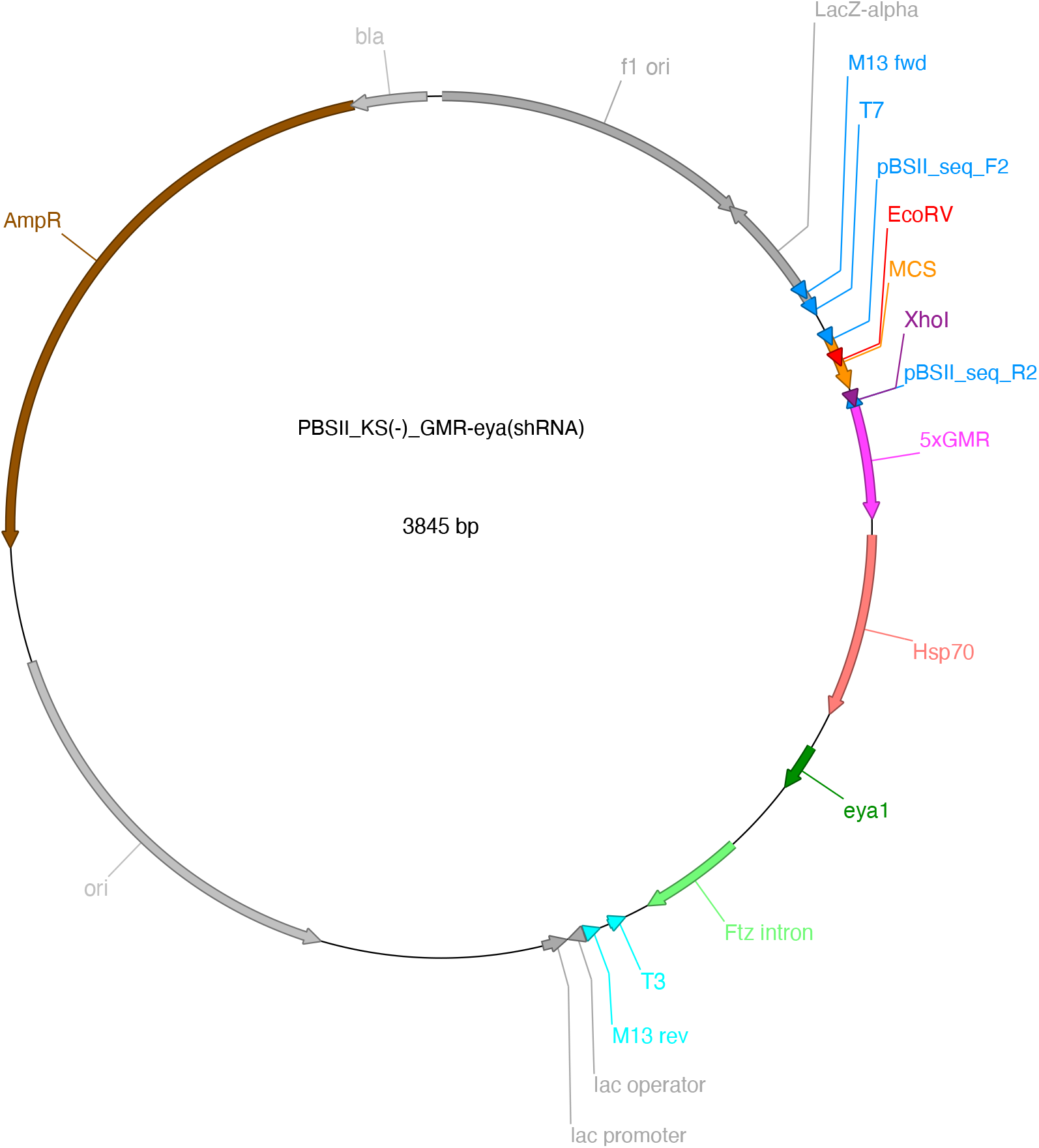
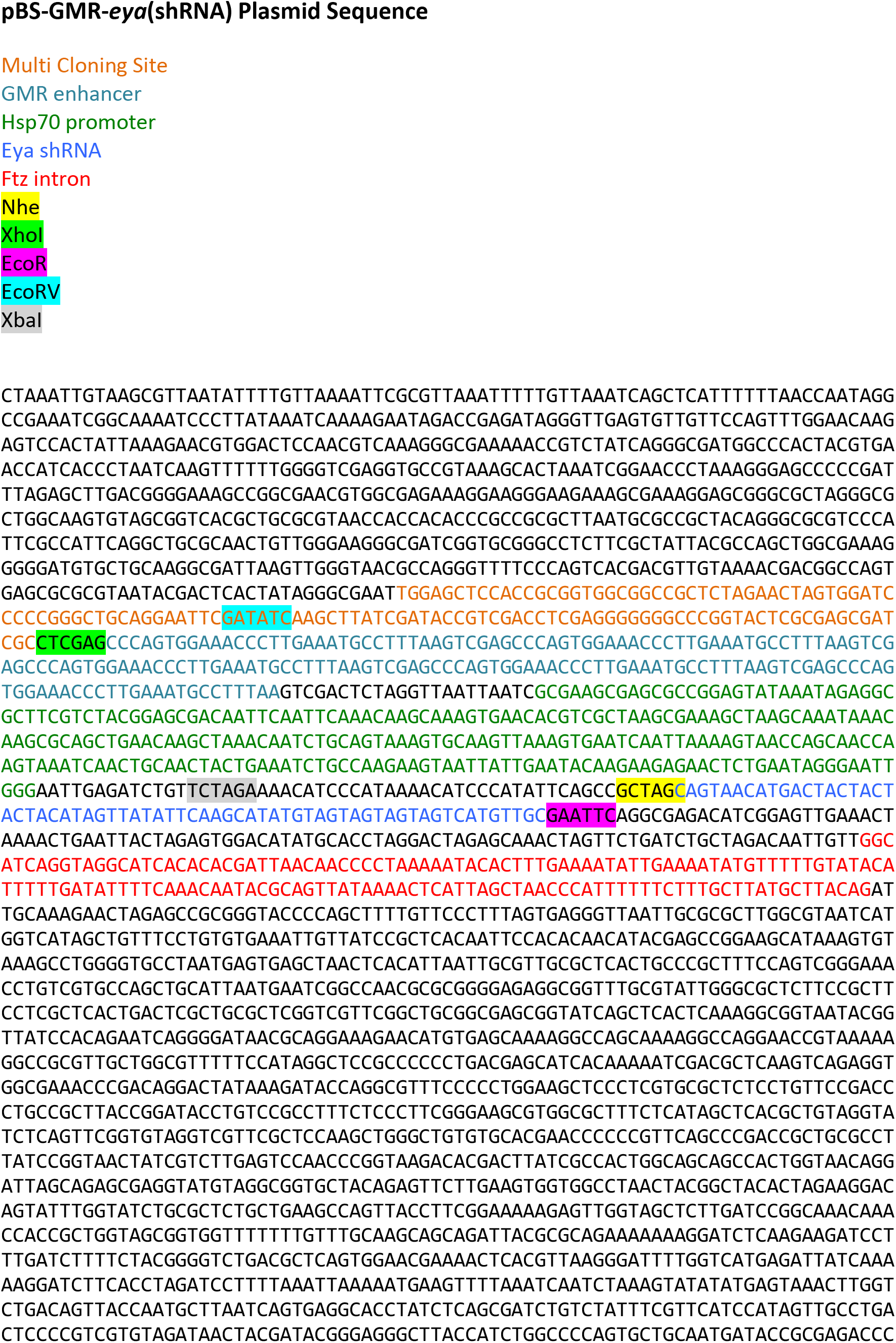
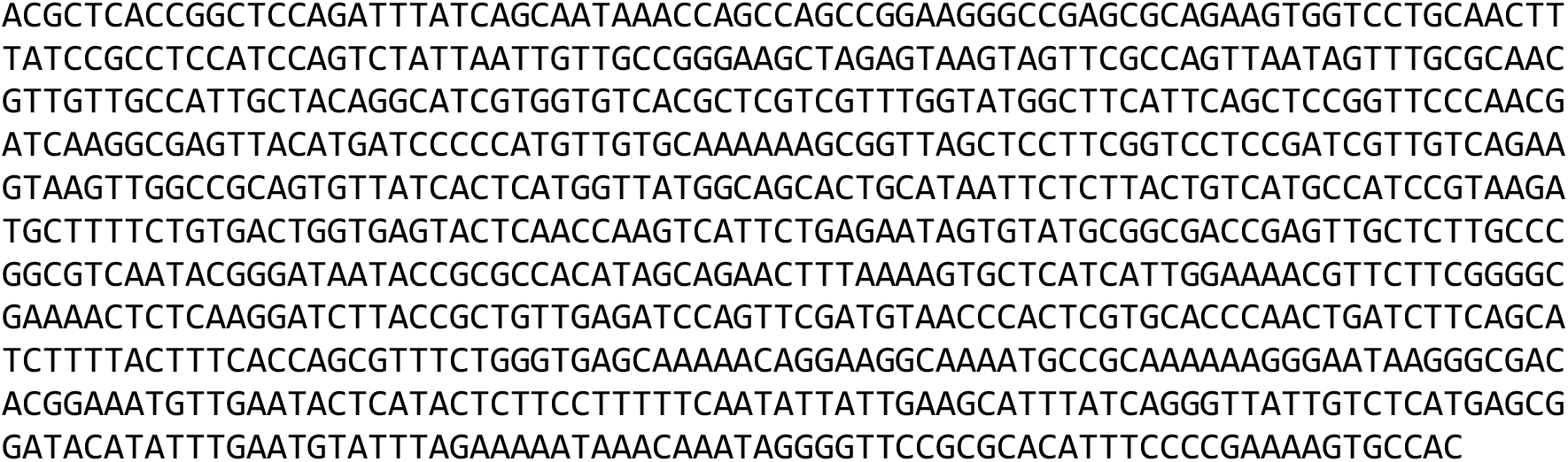
Nyberg et al.

## Supplementary Information

### Experimental Protocols

#### I. Selection of candidate sgRNAs to test

Candidate single-guide RNAs (sgRNAs) can be identified using flyCRISPR Optimal Target Finder (http://targetfinder.flycrispr.neuro.brown.edu, Gratz. et al. 2014). The sgRNA target site should be as close to the intended site of modification as possible. A length of 20 nucleotides works well, and an initial 5’ G or GG in the target site sequence is not necessary for in vitro transcription using T7 RNA polymerase. High stringency filtering is sufficient, and only NGG PAM sites should be utilized. Potential off-target sites should be minimized; zero predicted off-target sites is ideal.

The sgRNA target site sequence should be validated in the *Drosophila* strain or species genotype you plan to edit. This should be done by Sanger sequencing the putative site from the strain’s genomic DNA. Since sequence polymorphisms are prevalent across the genome of various stocks, the *Drosophila* reference genome sequence should only be taken as a guide, and the stock of interest should be sequence verified.

#### II. In vitro transcription (IVT) of sgRNA

##### Make the sgRNA DNA template for IVT

Perform a 50 μL PCR reaction using a proofreading polymerase (e.g. NEB Phusion HF DNA polymerase, New England Biolabs #M0530S) and the pU6-BbsI-chiRNA plasmid (Addgene #45946) as a template. Design PCR primers as follows to generate a ~120 bp PCR product:

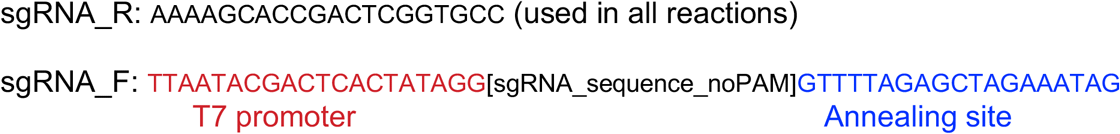

The T7 promoter sequence enables T7 RNA polymerase to initiate transcription. The Annealing site enables the primer to anneal to the plasmid template. Note that the PAM site is not included in the in vitro transcribed sgRNA.

Verify successful PCR amplification using an agarose gel - the product should be ~120 bp in length. Purify the PCR product using standard column purification (e.g. Qiaquick PCR Purification Kit, Qiagen #28106).

##### IVT

Use ~300 ng purified PCR DNA in a 20 μL MEGAscript in vitro transcription reaction (ThermoFisher #AM1333) supplemented with 0.5 μL Ribolock RNase inhibitor (ThermoFisher #EO0381). Incubate reaction at 37°C overnight in a thermocycler with a heated lid. Purify reaction products using an RNA cleanup column and elute in 20 μL nuclease-free dH_2_O (e.g. New England Biolabs Monarch RNA Cleanup Kit #T2040L). Successful IVT should yield >40 μg of RNA and should produce a large discrete band on an agarose gel.

#### III. Assembly of sgRNA-Cas9 RNPs

Mix together at room temperature to a final volume of 5 μL:

1.19 μL Cas9-NLS protein (IDT #1081058, 10 μg/μL)
0.38 μL 2M KCl
2.36 μg sgRNA
Nuclease-free dH_2_O to 5 μL

Incubate at room temperature for 10 min. Centrifuge in a microfuge at maximum speed for 10 min at room temperature. Transfer 4 μL of supernatant into new tube to be loaded into injection needles. Store at room temperature. Prepare RNPs fresh for each day of injections.

#### IV. Determination of sgRNA Cleavage Efficiency in Embryos

##### Embryo injections

Injections are performed in pre-cellularized embryos without dechorionation using Gompel and Schröder’s method (http://gompel.org/wp-content/uploads/2015/12/Drosophila-transformation-with-chorion.pdf). Injection of 35-40 embryos per batch of sgRNA RNPs should be sufficient. Also perform injections of 35-40 embryos with Cas9-NLS protein only or mock-injections as a control. After injection, rupture any embryos that were skipped during injection due to age or other defects with a needle. Then remove as much of the halocarbon oil as possible from the coverslip. Place coverslip with embryos on an egg-laying plate. Keep plate in a humid chamber at 25°C overnight.

##### Genomic DNA extraction

Injected embryos should be harvested ~24 hours post-injection. Use L1 larvae (preferable) or late-stage embryos, indicating survival of the injection process. Use the following single-fly genomic protocol (courtesy of Justin Kumar). Eight or more individuals should be sufficient to screen one sgRNA.

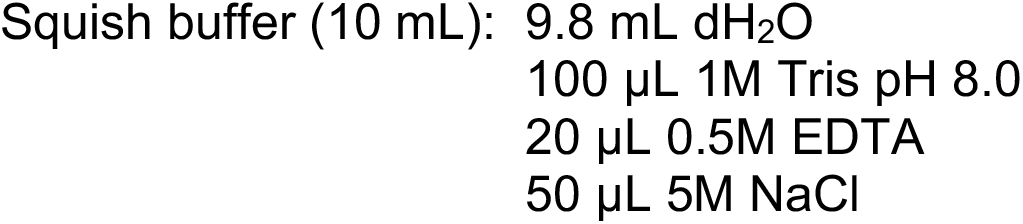

Before squishing, add 1 μL proteinase K (20 mg/mL) for every 100 μL of squish buffer.

1. Pick a single embryo/L1 larva with a pipet tip and grind it in 20 μL squish mix in a PCR tube. Pipette the mix several times. It is easy to lose the animal, so check under a dissecting scope to ensure that it is still inside the tube and ruptured (easier with larvae).
2. Incubate at 37°C for 30 min.
3. Incubate at 95°C for 5 min.
4. Store genomic preps at 4°C.

##### T7 endonuclease I assay

The DNA substrate for the T7EI digestion should be ~700-1200nt in length. This is made by PCR from the genomic DNA of a single embryo. The sgRNA target site should be located close to the center of the amplicon. Verify before running the T7EI assay that you can obtain a single robust PCR product and that T7EI digestion does not produce bands that overlap with predicted cleavage products generated by a sgRNA. Even PCR substrates from uninjected control embryos can produce multiple faint digestion products after T7EI digestion, presumably due to inherent sequence heterozygosity in the fly strain.

1. *Synthesize the T7EI DNA substrate*: Perform a 50 μL PCR reaction for each individual L1/embryo using 3-5 μL genomic DNA as template.
2. *Generate heteroduplexes*: Take 10 μL of PCR reaction in step 1 and perform the following denaturation and re-annealing in a PCR machine:

95°C – 3 min.
94°C – 1 min.
93°C – 1 min.
92°C – 1 min.
… continue downwards in 1 degree increments to…
4°C – 1min.
6°C – 10 sec.
8°C – 10 sec.
10°C – 10 sec.
12°C – hold
3. *Perform T7EI digestion*: To 10 μL re-annealed PCR DNA, add:
  2 μL 10X NEBuffer 2
  0.2 μL T7 Endonuclease I (New England Biolabs #M0302L)
  7.8 μL dH_2_O.
  Incubate at 37°C for 1 hour.
4. *Analysis of reaction*: Run in a 2% (w/v) agarose gel: 10 μL undigested PCR product side-by-side with 20 μL T7EI reaction products. Ethidium bromide and 0.5X TBE should be used to increase sensitivity to see faint digestion products. Samples with cleavage products at expected sizes from RNP-injected animals that are not present in mock-injected controls are indicative of sgRNA-guided cleavage. Bands may be very faint. It is not unusual for > 50% of individuals to have cleavage products for a good sgRNA.

#### V. Donor plasmid design

The CRISPR/Cas9 system can be used to introduce various modifications (e.g. protein tags, precise mutations) into the genome of *Drosophila* via homology-directed repair (HDR). To do so, a donor plasmid carrying the intended modification must be introduced into the embryo along with a sgRNA and Cas9-NLS.

The first step is to computationally assemble the donor plasmid using an informatics tool such as Benchling (https://www.benchling.com). The donor plasmid typically consists of 5 pieces:

1. backbone plasmid with a negative selection marker
2. modification of interest
3. positive transformation marker
4. left homology arm
5. right homology arm

##### (1) Backbone plasmid with negative selection marker

All the pieces necessary for genome editing via HDR need to be inserted via Gibson assembly into a plasmid with a negative selection marker that can be used to screen against integration of the entire plasmid into the genome. We use pBS-GMR-*eya*(shRNA), described in the paper. It carries a short hairpin RNAi agent against *eya* mRNA transcripts. Its transcription is driven by the eye-specific GMR enhancer. If it integrates into the *Drosophila* genome, it results in small eyes and can be used in any line with normal eye morphology. The pBS-GMR-*eya*(shRNA) plasmid is 3,845 bp and its annotated sequence is in Supplementary File 1.

The plasmid is linearized via restriction enzyme digestion before Gibson assembly. pBS-GMR-*eya*(shRNA) can be linearized with *EcoRV* (recognition sequence: GATATC), which is located in the multi-cloning site of pBluescript. Since *EcoRV* digestion generates blunt ended fragments, no nucleotides will be removed by the 5’->3‘ exonuclease activity of the Gibson assembly, and thus the assembled insert should be placed right at the cut site.

##### (2) Modification of interest

The modification of interest (MOI) should be placed as close to the sgRNA target site as possible to minimize the possibility of homologous recombination occurring between the sgRNA target site and the MOI. We have had success with the MOI located less than 30 bp from the sgRNA target site. Placement within the sgRNA site is ideal, as it will also inactivate the sgRNA site in the donor plasmid.

##### (3) Positive transformation marker

The 3xP3-DsRed marker gene is used to screen for positive integration of the intended modification via HDR. Note that 3xP3-DsRed fluorescence is only visible in a small number of ommatidia in a wildtype eye color background, making the fluorescence difficult though not impossible to observe.

The 3xP3-DsRed cassette from the pScarlessHD-DsRed plasmid (Addgene #64703) is flanked by piggyBac transposition sites (TTAA) that can be used to cleanly excise the entire marker gene after successful integration of the modification via HDR. After excision, the remaining genome sequence will be reduced to a single TTAA site. Thus, the scarless cassette should be placed either in a native TTAA site near the sgRNA site (ideally less than 30 bp) or within a TTAA site in the intended modification. Placement within a TTAA inside the sgRNA site is ideal, as it will also inactivate the sgRNA site in the donor plasmid.

##### (4) Left and right homology arms

For successful HDR, homology arms of native genomic sequence must be present on either side of the MOI and sgRNA target site. Lengths of ~1000 bp are standard. Lengths can be slightly increased or reduced to provide ideal sequences for Gibson assembly (e.g. moderate GC content and nonrepetitive sequence).

###### Important

**If you cannot inactivate the sgRNA site in the donor plasmid either by inserting the MOI or scarless-DsRed cassette into the sgRNA core, then you need to mutate at least a single basepair in the PAM site or the sgRNA core of the donor plasmid.**

##### Design of primers for Gibson assembly

Once an ideal donor plasmid is computationally designed, Gibson assembly can be used to assemble the necessary DNA fragments into the donor plasmid. DNA fragments can be generated in 3 ways:

1. PCR of genomic or plasmid template
2. Restriction digest of plasmid
3. Commercial de novo synthesis (e.g. IDT GBlocks)

In most cases, the backbone plasmid is generated via *EcoRV* digest, and the homology arms are PCR amplified from genomic DNA from the same *Drosophila* strain or species to be used for injections. The positive transformation marker is typically generated via PCR from pScarlessHD-DsRed plasmid DNA. The modification of interest can be generated either via PCR or de novo synthesis.

To design ideal primers to generate DNA fragments for Gibson assembly, use the NEBuilder tool http://nebuilder.neb.com with the following build settings:

Product Kit: NEBuilder HiFi DNA Assembly Master Mix
Minimum Overlap: 30 nt
Circularize: Yes
PCR Polymerase/Kit: Phusion High-Fidelity DNA Polymerase (HF Buffer)
PCR Primer Conc.: 500 nM
Min. Primer Length: 18
Max. Primer Length: 60 (not a build setting, but necessary for standard IDT order)

Try to alleviate flagged issues if possible, though not all issues can be resolved. For example, you cannot change the ends of the cut backbone plasmid, even if they are not ideal for Gibson assembly. Ends of homology arms can be slightly altered to improve Gibson overlap regions, and a synthesized gBlocks fragment can be altered to do the same.

Avoid placing repetitive regions like the very ends of the piggyBac transposition sites into Gibson overlap regions. Overlap regions can be slightly altered via junction properties in NEBuilder. For the scarless 3xP3-DsRed cassette, the 17 bp at both ends of the cassette are identical (5’-TTAACCCTAGAAAGATA-3’) and thus should not be used in Gibson overlap regions. If synthesizing the MOI via gBlocks, one potential workaround is to extend the gBlocks fragment through the adjacent transposon end of the 3xP3-DsRed cassette to place the Gibson overlap region deeper into a nonrepetitive region of the cassette. We have verified that the following sequences within the 3xP3-DsRed cassette can be used as Gibson overlap regions:

piggyBac left (5’) region: 5’-GTCGTTATAGTTCAAAATCAGTGACACTTA-3’
piggyBac right (3’) region: 5’-AGATAATCATGCGTAAAATTGACGCATGTG-3’

Once all primers are designed, verify that they all will bind in your computationally assembled donor plasmid.

#### VI. Construction of the donor plasmid via Gibson assembly

##### (1) Backbone plasmid with negative selection marker

Digest 5-10 μg of pBS-GMR-*eya*(shRNA) with EcoRV-HF (New England Biolabs #R3195S) at 37°C for 15 min. Digested product should be run on a 1% agarose gel, using multiple lanes to accommodate the large volume of digest. Bands of linearized plasmid should be quickly and carefully excised from gel, minimizing exposure to UV light, and purified using Monarch DNA Gel Extraction Kit (New England Biolabs #T1020S) to avoid contamination of Gibson assembly reactions with trace uncut plasmid.

##### (2) Scarless DsRed cassette

Perform a 50 μL PCR reaction using a proofreading polymerase (e.g. New England Biolabs Phusion HF) and 30-50 ng of pScarlessHD-DsRed plasmid as template (Addgene #64703). Touchdown PCR is recommended to reduce nonspecific bands. The entire PCR product should be run on a 1% agarose gel, and the desired product should be gel extracted as above to avoid contamination of the Gibson assembly reaction with template plasmid. Similarly, gel extraction should be performed on any other PCR reaction that uses plasmid as template.

##### (3) Left and right homology arms

Perform a 50 μL touchdown PCR reaction for each homology arm using a proofreading polymerase (e.g. New England Biolabs Phusion HF) and 50 ng of genomic DNA from the same *Drosophila* strain or species that will be used for injections. If multiple bands are present, then purify via gel extraction. Otherwise, standard column purification is sufficient (e.g. Qiaquick PCR cleanup, Qiagen #28106).

##### (4) Synthesized DNA fragments

Any synthesized DNA fragment (e.g. IDT GBlocks) should be briefly centrifuged and resuspended in molecular grade dH_2_O to a final concentration of 10 ng/μL. Incubate at 50°C for 15 minutes to facilitate better resuspension.

##### Assembly

Empirically determine concentration of all fragments using a fluorometer (e.g. Qubit) or spectrophotometer (e.g. Nanodrop). For a 5 piece Gibson assembly reaction, fragments should be added in equimolar amounts, with total DNA content of the reaction not exceeding 0.5 pmol. The combined volume of DNA fragments should be 10 μL or less. Adjust volume using dH_2_O. 0.08 - 0.1 pmol per fragment works well. The accompanying Gibson assembly calculator can be used to determine appropriate volumes.

To perform the Gibson assembly reaction:

1. Mix all DNA fragments together. Combined volume should be less than 10 μL. Add dH_2_O to 10 μL.
2. Mix 10 μL of combined DNA fragments with 10 μL NEBuilder HiFi DNA Master Mix (New England Biolabs #E2621). Mix well.
3. Incubate at 50°C for 1 hour in a thermocycler with heated lid.
4. Transform into competent *E. coli*. As even successful Gibson assembly reactions produce a small number of colonies, it is important to use *E. coli* with as high transformation efficiency as possible. Electrocompetent *E. coli* typically have higher efficiency than chemically competent *E. coli*.

A successful reaction will produce one to several hundred colonies. Performing a negative control reaction in parallel is useful to distinguish a successful low-yield reaction from non-specific colonies. Negative control reactions typically contain NEBuilder HiFi DNA Master Mix and only the backbone plasmid and scarless DsRed cassette fragments, as these are most likely to introduce contaminants. Individual colonies can be picked and screened via PCR for successful assembly across 1-2 junctions. Confirm correct assembly of the entire inserted region via Sanger sequencing. Polymorphisms in noncoding regions of homology arms are not uncommon, but ensure that there are no disabling mutations in the scarless DsRed cassette or coding regions of the homology arms.

##### Purification of the donor plasmid

Purify the donor plasmid DNA for injection using the HiSpeed Plasmid Midi Kit (Qiagen #12643) with additional removal of endotoxins using two reagents from the EndoFree Plasmid Mega Kit (Buffer ER and Buffer QN, Qiagen #12381) to reduce toxicity in injected embryos. The Midi Kit is used as directed by the manufacturer with several modifications, as indicated in red below:

1. Pellet 50 mL of an overnight LB culture at 6000 x g for 15 min at 4°C.
2. Decant supernatant and resuspend pellet in 6 mL Buffer P1 with added RNase A by vortexing.
3. Add 6 mL Buffer P2 and mix well by inverting 4-6 times. Incubate at RT for 5 min.
4. During incubation, screw the cap onto the outlet nozzle of the QIAfilter Cartridge. Place the cartridge into a rack or fresh 50 mL conical tube.
5. Add 6 mL prechilled Buffer P3 to lysate and mix well by inverting 4-6 times.
6. Pour lysate into the QIAfilter Cartridge and incubate at RT for 10 min.
7. Remove the cap, insert the plunger, and filter the solution through the syringe filter into a fresh 50 mL conical tube.
8. Add 1 mL (EndoFree Mega) Buffer ER to the filtered solution and incubate on ice for 30 min.
9. During incubation, equilibrate a HiSpeed Tip with 4 mL Buffer QBT.
10. Apply the incubated solution from step 8 to the QBT-equilibrated HiSpeed Tip and allow to flow through.
11. Wash the HiSpeed Tip 2 x 10 mL with Buffer QC.
12. Place the HiSpeed Tip over a fresh 50 mL conical tube and elute by applying 5 mL (EndoFree Mega) Buffer QN.
13. Add 3.5 mL isopropanol to the eluted solution. Mix by inverting and incubate at RT for 5 min.
14. During incubation, remove the plunger from a 20 mL syringe and attach the QIAprecipitator Module onto the outlet nozzle.
15. Place the QIAprecipitator over a spare 50 mL conical tube. Transfer the eluate mixture into the syringe and insert the plunger. Filter the mixture through using constant pressure.
16. Remove the QIAprecipitator from the syringe and pull out the plunger. Reattach the QIAprecipitator and add 2 mL 70% EtOH to the syringe. Insert the plunger and push the 70% EtOH through.
17. Remove the QIAprecipitator from the syringe and pull out the plunger. Attach the QIAprecipitator again and insert the plunger. Dry the membrane by pressing air through the QIAprecipitator. Repeat this step several times.
18. Dry the outlet nozzle of the QIAprecipitator with a Kimwipe.
19. Remove the plunger from a new 5 mL syringe, attach the QIAprecipitator and hold the outlet over a 1.5 mL collection tube. Add 1 mL Buffer TE to the syringe. Insert the plunger and elute the DNA into the collection tube using constant pressure.
20. Remove the QIAprecipitator from the 5 mL syringe and pull out the plunger. Re-attach the QIAprecipitator to the syringe.
21. Transfer the eluate from step 19 to the 5 mL syringe and elute for a second time into the same 1.5 mL tube.

This final elution should be performed using TE buffer to maximize recovery of the plasmid DNA. However, TE buffer is not appropriate for injections, and the donor plasmid needs to be concentrated before injection. Perform an ethanol precipitation as follows:

1. Estimate volume of DNA solution and add 1/10 volume of 3M sodium acetate pH 5.2. Mix well.
2. Add 3 volumes of 100% molecular-grade ethanol.
3. Incubate at −80°C for 30 minutes.
4. Spin at max speed for 15 minutes at 4°C. Split into multiple 1.5 mL tubes if necessary.
5. Remove supernatant and wash pellet twice in 800 μL of 70% ethanol.
6. After final wash, remove supernatant and allow to air-dry at RT 5-10 minutes.
7. Resuspend in 40 μL nuclease-free dH_2_O.
8. Measure concentration using NanoDrop or Qubit. Final concentration should be ~240 nM or higher.

#### VII. Injection and screening of transformants

Injections are performed in pre-cellularized embryos without dechorionation using Gompel and Schröder’s method (http://gompel.org/wp-content/uploads/2015/12/Drosophila-transformation-with-chorion.pdf). Injections should be performed in the same *Drosophila* strain or species used for sgRNA prescreening.

Mix together at room temperature:

1.19 μL Cas9-NLS protein (IDT #1081058, 10 μg/μL)
0.38 μL 2M KCl
2.36 μg IVT sgRNA
0.60 pmoles donor plasmid DNA
Nuclease-free dH_2_O to 5 μL final volume

Incubate at room temperature for 10 min. Centrifuge in a microfuge at maximum speed for 10 min at room temperature. Transfer 4 μL of supernatant into new tube to be loaded into injection needles. Store at room temperature. Prepare RNPs fresh for each day of injections.

Injection of 300-350 embryos is typically sufficient to obtain at least 1 germline transformant. After injection, remove as much oil as possible and place coverslip with injected G0 embryos in a standard food vial. Keep the vial in a humid chamber at 25°C overnight.

Once G0 adults eclose, they should be individually crossed to healthy virgins or males from a wild-type or appropriate balancer line. We typically use *w^1118^* flies for injections and the initial cross in order to maintain a consistent genetic background. G1 adults are screened for the expression of 3xP3-DsRed and the absence of *eya*(*shRNA*) phenotypes. Positive G1 adults typically contain the desired edit and should be individually crossed to an appropriate balancer. Note that you might obtain multiple positive G1 adults from the same G0 parent. These may or may not be independent genome modifications. However, you can be confident that G1 adults taken from different G0 parents will have independent edits.

Once lines are established and stable, verification of the anticipated editing/modification needs to be done by PCR analysis and Sanger sequencing. Errors do occur.

#### VIII. Removal of the DsRed marker with piggyBac transposase

To precisely excise the 3xP3-DsRed marker cassette and achieve scarless genome editing, set up the following crosses:

(P) Cross DsRed+ lines to flies expressing the piggyBac transposase. Bloomington stock 32070 contains a piggyBac transposase transgene under control of the *α-tubulin* promoter and tightly linked to a 3XP3-CFP transgenic marker. This is located on chromosome 2. The stock also contains 3rd chromosome balancers (MKRS/TM6B,Tb), facilitating tracking of the 3rd chromosomes independent of the piggyBac transposase. If DsRed is on the X chromosome, cross virgin DsRed+ females to males of the piggyBac transposase line.
(F_1_) If your 3xP3-DsRed gene is on the X chromosome, select several DsRed+/CFP+ males. If your 3xP3-DsRed gene is on an autosome, select several DsRed+/CFP+ males. Cross males to 10-20 virgin females with an appropriate balancer. The piggyBac transposase is only weakly efficient, so DsRed+ should still be visible albeit mosaic in F1 flies.
(F_2_) If the DsRed was on an autosome, select single flies that have the appropriate balancer chromosome and are both DsRed- and CFP- and cross again to an appropriate balancer to make a balanced stock. If the DsRed was on the X chromosome, select single female flies that are both DsRed- and CFP-, and cross to males from an appropriate balancer line to make a balanced stock. Removal of DsRed typically occurs 10% of the time or less, so make sure crosses are large enough to produce hundreds of F2 progeny to screen through.

If the genome editing has been performed on a species other than *D. melanogaster*, it will be necessary to inject the 3xP3-DsRed lines with a plasmid vector containing the piggyBac transposase gene under alpha-tubulin promoter control. This plasmid is commercially available (Drosophila Genome Resources Center #1155). Injections can be performed in pre-cellularized embryos without dechorionation using Gompel and Schröder’s method (http://gompel.org/wp-content/uploads/2015/12/Drosophila-transformation-with-chorion.pdf). Injections should be performed using a concentration of 0.6 mg/mL plasmid DNA dissolved in 0.1 mM Sodium Phosphate pH 7.8 + 5 mM KCl. Cross individual G0 adults to an appropriate strain and screen G1 adult offspring for the absence of DsRed eye fluorescence. Since the pBac transposase plasmid vector requires active P element transposase to integrate into an injected embryo’s genome, there should be no retention of the transposase gene in G1 adults.

To ensure that the genome edit is still present after scarless excision, verify via PCR analysis and Sanger sequencing.

